# Forelimb motor recovery by modulating extrinsic and intrinsic signaling as well as neuronal activity after the cervical spinal cord injury

**DOI:** 10.1101/2024.06.22.600167

**Authors:** Hirohide Takatani, Naoki Fujita, Fumiyasu Imai, Yutaka Yoshida

## Abstract

Singular strategies for promoting axon regeneration and motor recovery after spinal cord injury (SCI) have been attempted with limited success. Here, we propose the combinatorial approach of deleting extrinsic and intrinsic factors paired with neural stimulation, will enhance adaptive axonal growth and motor recovery after SCI. We previously showed the deletion of *RhoA* and *Pten* in corticospinal neurons inhibits axon dieback and promotes axon sprouting after lumbar SCI. Here, we examined the effects of *RhoA;Pten* deletion coupled with neural stimulation after cervical SCI. This combinatorial approach promoted more boutons on injured corticospinal neurons in the spinal cord compared to sole *RhoA*;*Pten* deletion. Although *RhoA*;*Pten* deletion does not promote motor recovery in the forelimb after SCI, stimulating corticospinal neurons in those mice results in partial motor recovery. These results demonstrate that a combinatorial approach that pairs genetic modifications with neuronal stimulation can promote axon sprouting and motor recovery following SCI.

## INTRODUCTION

Recovery of motor function after spinal cord injury (SCI) is likely to require substantial neural rewiring. In the adult central nervous system, full recovery from SCI is rarely observed. The corticospinal tract (CST), the major descending circuit that controls voluntary movements, conveys motor commands from the sensorimotor cortex to the spinal cord making it an important target for axon regeneration following SCI^1–3^. Although numerous potential therapeutic methods have been attempted, no single approach has fully reconstructed the injured CST or recovered motor function after SCI. We propose that a combinatorial strategy targeting signaling pathways and neural activity may yield superior results compared to individual therapies.

Several molecules, such as phosphatase and tensin homolog (Pten), KLF7, Rho, and Sox11, have been shown to prevent axon regeneration after SCI ^4–7^. Pten is an enzyme that suppresses the mechanistic target of rapamycin (mTOR; a critical signaling component for axon growth) which prevents new protein synthesis and neuronal growth ^8^. Deletion of *Pten* promotes significant regeneration of CST axons after SCI, facilitating robust extension of axons across the lesion site ^4^. The Rho family of small GTPases, including RhoA, are important regulatory signaling molecules that are triggered by extrinsic molecules, such as myelin-associated inhibitors, repulsive axon guidance molecules, and inhibitory extracellular matrix molecules after SCI ^7^. The Rho signaling pathway is involved in actin cytoskeletal dynamics that can lead to growth cone collapse and axon growth inhibition. Pharmacological Rho inhibitors enhance sprouting and regeneration of corticospinal (CS) axons after SCI, but the extent of regeneration is variable ^7, 9–11^. We previously showed that co-deletion of *RhoA* and *Pten* from sensorimotor cortex prior to SCI resulted in reduced CS axonal dieback and enhanced rewiring of CS circuits to cerebral cortex, spinal cord, and hindlimb motor units; however, this approach to changing the internal growth state of corticospinal neurons (CSNs) alone did not promote axon regrowth through the lesion and did not enhance hindlimb motor recovery after SCI^12^. A method for eliciting and strengthening novel CS circuits may be required for capitalizing on the enhanced axonal plasticity resulting from *RhoA* and *Pten* deletion.

Neuronal stimulation is a robust approach to drive CS circuit plasticity that may promote motor recovery. Previous studies indicate that electrical stimulation of the cortex can transform neuronal circuits from a nonfunctional to a highly functional state and promote extensive sprouting of CSNs which restores neurological function after SCI ^13, 14^. In another study, chemogenetic stimulation of spinal neurons using designer receptors activated only by designer drugs (DREADDs), can induce axon sprouting and restore locomotion after SCI by administering actuator ligands ^15^. In addition, such chemogenetic stimulation of CSNs in both supraspinal centers and spinal relay stations results in enhancement of neuronal rewiring and locomotor recovery ^16^.

Together, neuronal stimulation of the *RhoA*;*Pten* deleted axons, which have regained the potential to grow and sprout, is a novel strategy to enhance axon regeneration and motor recovery after SCI.

In this study, we tested the hypothesis that combining *RhoA*;*Pten* deletion in CSNs with neuronal stimulation synergistically enhances rewiring of CS circuit and forelimb functional recovery after cervical SCI in mice. Our results demonstrated that this combinatorial intervention spurred CS axon regrowth and adaptive rewiring, plus achieved partial restoration of forelimb motor recovery. Our results suggest that combinatorial treatment strategies may ultimately lead to a multi-pronged treatment regimen that maximizes motor recovery in individuals impacted by traumatic SCI.

## RESULTS

A recent study examining recovery after SCI showed that modulation of extrinsic and intrinsic signaling pathways through deletion of both *RhoA* and *Pten* suppresses axon dieback at the thoracic level of the spinal cord ^12^. Excitation of CSNs has also been shown to promote axon sprouting following CST injury ^14^. We hypothesized that combining these approaches would augment recovery in the spinal cord following acute trauma. We tested this two-pronged approach in mice by deleting *Pten* and *RhoA* in CSNs to modulate both intrinsic signaling and the response to extrinsic inhibitory pathways, respectively, and combining those genetic approaches with CSN activation using chemogenetics (Fig. 1). We determined the functional and anatomical effects of this unique combinatorial approach on CSN circuit recovery after cervical SCI.

**Figure 1.**
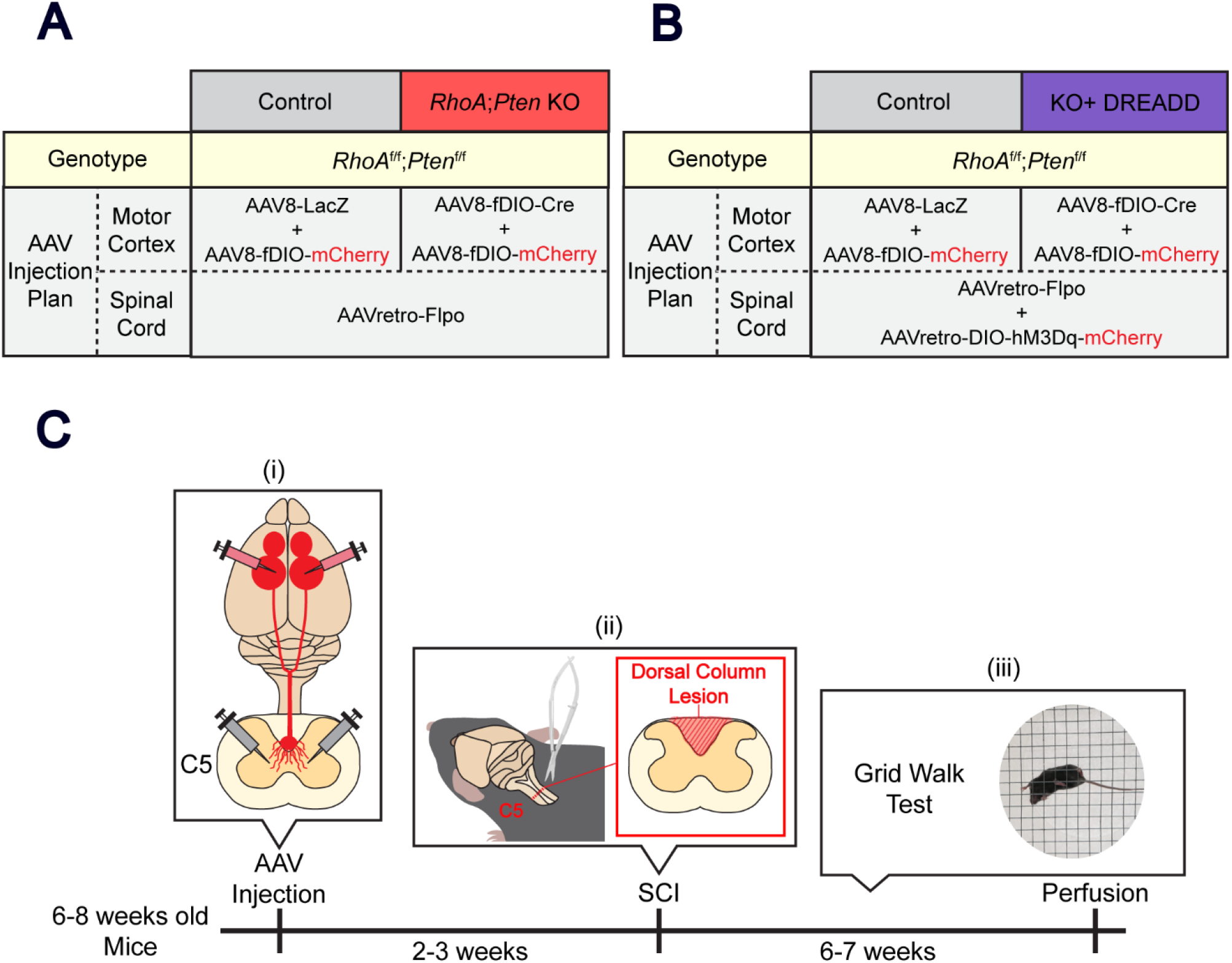
Experimental procedure. (A-B) AAV injection strategy. RhoA;Pten KO mice were generated by injecting AAVs to induce RhoA and Pten KO, and mCherry expression in CST axons. KO+DREADD mice were generated by injecting RhoA;Pten KO mice with AAVs for hM3Dq and mCherry expression. Control mice were only injected with AAVs for mCherry expression. (C) Experimental schematic and timeline. (i) AAVs were injected into the forelimb area of the sensorimotor cortex and the C5 level of the spinal cord in 6-8 week old RhoAf/f;Ptenf/f mice. (ii) The C5 dorsal column lesion was performed 2-3 weeks after AAV injections and (iii) the grid walking test was performed weekly thereafter followed by perfusion at 6-7 weeks post-injury.

### Cre recombinase induces sufficient *RhoA*;*Pten* deletion

We first determined the effects of genetic deletion of *RhoA* and *Pten* in CSNs injecting AAV- Cre into the sensorimotor cortex (Fig. 2). By co-injection of AAV-DIO-GFP into the sensorimotor cortex, the injection of GFP^+^ Cre showed lower expression of both RhoA and Pten proteins compared to the control (Fig. 2B). Quantitative analysis also showed significantly lower protein levels in the GFP^+^ Cre, indicating that the Cre expression induced sufficient genetic co-deletion of *RhoA* and *Pten* (Fig. 2C).

**Figure 2.**
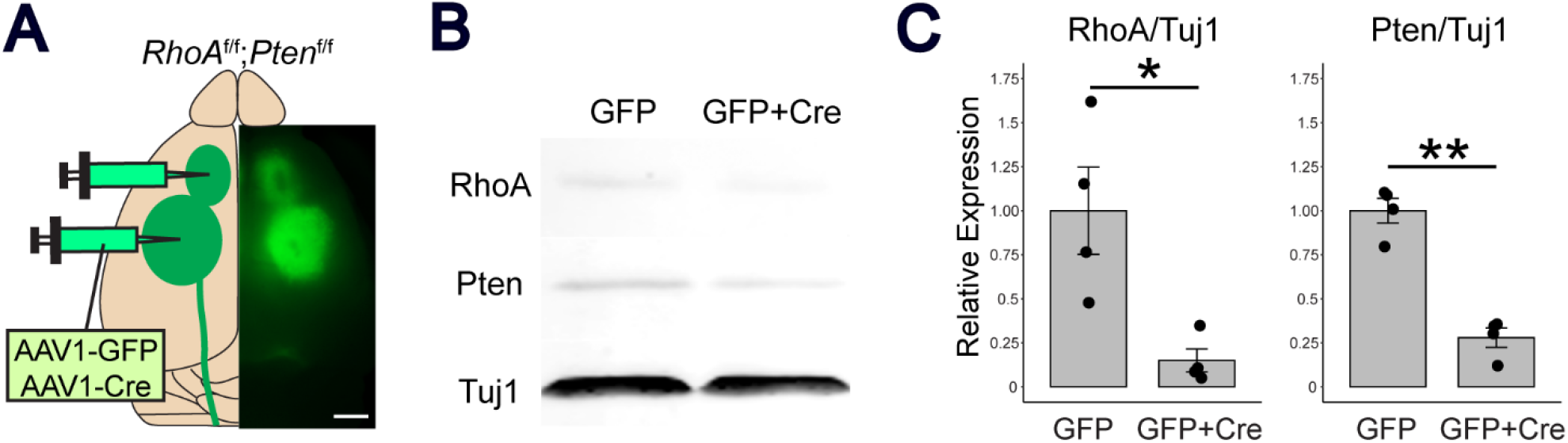
Genetic deletion of RhoA;Pten in the sensorimotor cortex. (A) Representative brain illustration showing the AAV1-Cre and AAV1-GFP injection strategy. AAVs were injected into the rostral area (RFA)and caudal forelimb area (CFA) of RhoAf/f;Ptenf/f mice. Photographic image on the right shows GFP expression in the cerebral cortex. Scale bar, 2 mm. B, Western blot of RhoA and Pten proteins at the AAV injection site of the cerebral cortex in RhoAf/f;Ptenf/f mice 2 weeks after AAV injections. C, Quantitative analysis of RhoA and Pten protein band intensity from Western blots for GFP mice (n = 4) and GFP+Cre mice (n = 4). Unpaired t-test, *p<0.05, **p<0.005.

### Combination of *RhoA;Pten* deletion in CSNs and CSN activation suppresses axon dieback after SCI

To activate CSNs, we used hM3Dq, an excitatory version of a designer receptor exclusively activated by designer drugs (DREADDs). Activation of hM3Dq by addition of the ligand, deschloroclozapine (DCZ), was evaluated by examining cFos expression in control and KO (*RhoA; Pten* deletion)+DREADD mice. The number of c-Fos^+^ and mCherry^+^ CSNs in KO+DREADD mice was significantly higher than the control, indicating that Cre induced the excitatory DREADD and DCZ functioned as an actuator ligand (Fig. 3).

**Figure 3.**
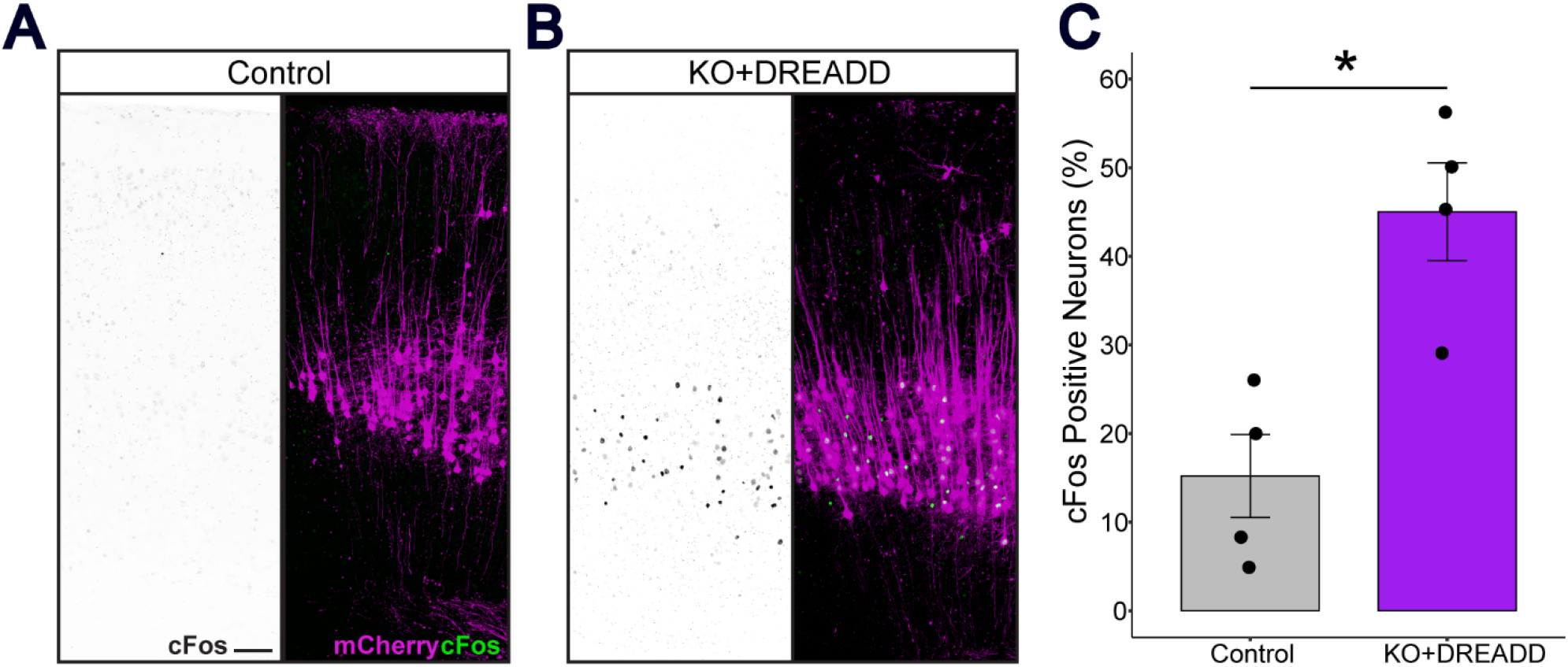
Neuronal stimulation by excitatory DREADD in the sensorimotor cortex induced cFos expression. (A-B) Representative images of Control mice and KO+DREADD mice. Left, cFos (black); right, cFos (green), mCherry (magenta). Scale bar, 100 µm. (C) Quantitative analysis of cFos+ mCherry neurons in Control mice (n = 4) and KO+DREADD mice (n = 4). Unpaired t-test, *p<0.05.

Next, sagittal sections of the spinal cord were examined to evaluate axon dieback in the CST (Fig. 4). In the two control groups, more than 50% of axons were present at 200-500 µm rostral to the lesion in the dorsal funiculus, indicating axon dieback (Fig. 4A, C). In contrast, more than 50% of axons were observed at 100 µm rostral to the lesion in KO and KO+DREADD mice (Fig. 4B, D). The CST axon indices in KO and KO+DREADD mice showed statistically significant less axon dieback compared to their corresponding controls, indicating that deletion of *RhoA* and *Pten* suppressed axon dieback (Fig. 4E-F). To further evaluate the differences between the KO and KO+DREADD mice, we measured the improvement in axon dieback suppression over the control. In the KO+DREADD mice, the axon dieback suppression ratio was significantly higher at 200 µm rostral to the lesion in KO+DREADD mice compared to KO alone (Fig. 4F).

**Figure 4.**
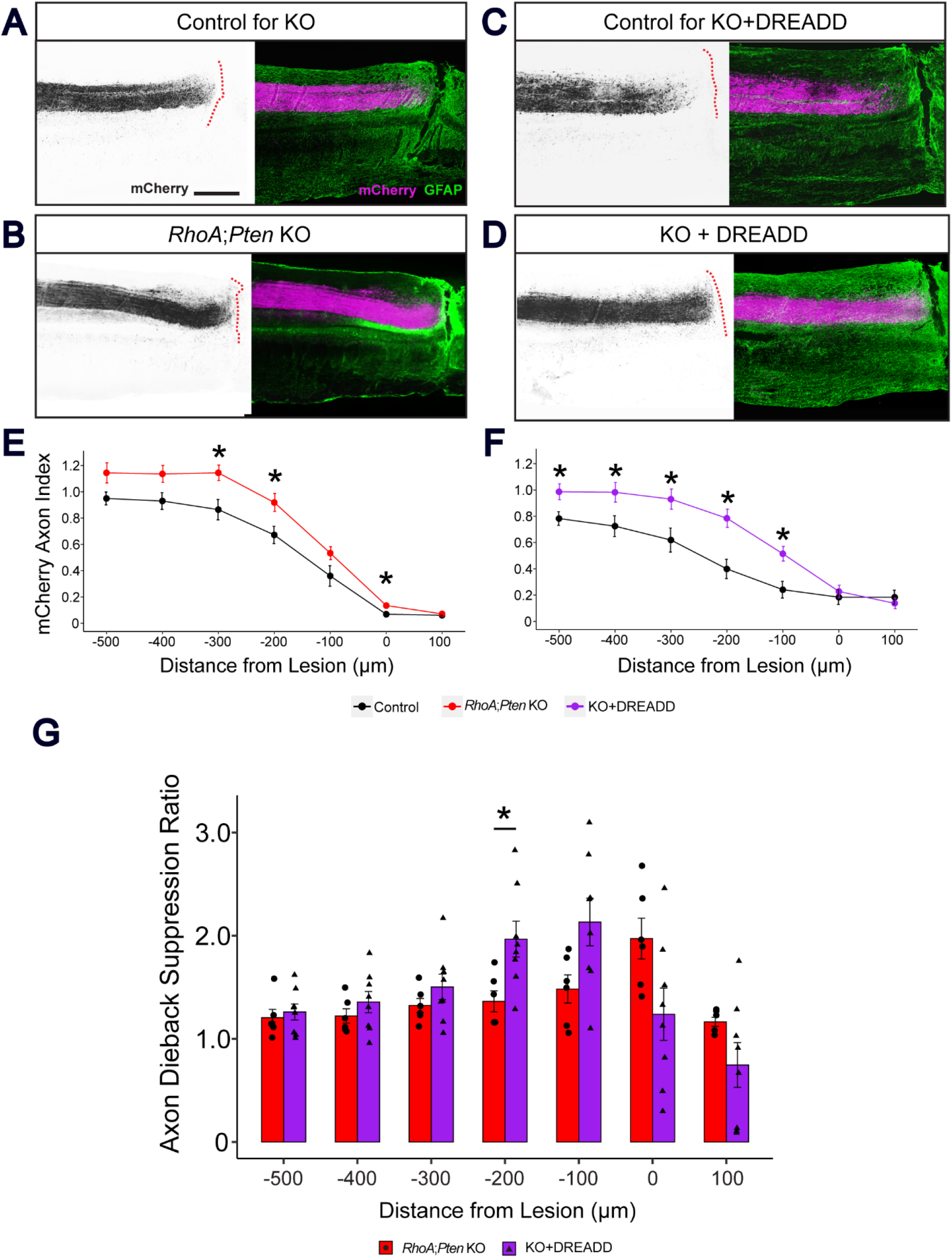
RhoA;Pten KO+DREADD mice show greater suppression of axon dieback in the CST than RhoA;Pten KO mice after SCI. (A-D) Representative images of mCherry+ CST axons in the cervical spinal cord rostral to the lesion in KO control mice (A), RhoA;Pten KO mice (B), KO+DREADD control mice (C) and KO+DREADD mice (D). Sagittal view of the cervical spinal cord at 42 DPI (A-B) and 49 DPI (C-D). Dotted red lines indicate the border of the GFAP+ glial scar and the GFAP- fibrotic scar. Left, mCherry (black); right, GFAP (green) and mCherry (magenta). Scale bar, 500 µm. (E) Quantification of CST axons in the dorsal funiculus in KO control mice (n = 6), RhoA;Pten KO mice (n = 7), KO+DREADD control mice (n = 6) and KO+DREADD mice (n = 8). Wilcoxon rank sum exact test, *p<0.05. (F) Improvement rates of RhoA;Pten KO and KO+DREADD mice. Wilcoxon rank sum exact test, *p<0.05.

These results suggest that *RhoA;Pten* deletion suppresses axon dieback at cervical levels, similar to those observed in the lumbar level of the spinal cord. Moreover, the combination of excitatory DREADD and *RhoA; Pten* deletion resulted in less axon dieback than *RhoA;Pten* deletion alone.

### Combination of *RhoA;Pten* deletion and excitatory DREADD promotes forelimb motor recovery

To determine whether the combination of *RhoA;Pten* deletion in CSNs with activation promotes enhanced motor recovery after SCI, we subjected mice to a grid-walking test. In this test of skilled locomotor behavior, the slip rate index was higher in the left forepaw in KO mice compared to their controls whereas no difference was observed for the right forepaw which led to higher value in KO mice for both forepaws (Fig. 5A). In contrast, the KO+DREADD mice showed significantly lower slip rates at 21-35 days post-injury (DPI) in both forepaws (Fig. 5B). Left forepaw slip rate indices (a normalized value, each rate divided by the pre-injury rate) in the KO+DREADD mice were significantly lower than those of the KO mice at 21 and 28 DPI (Fig. 5C). Similarly, right forepaw slip rate indices were found to be significantly lower for the KO+DREADD mice at 21 and 35 DPI. When both forepaws were examined together, the slip rate was significantly lower for the KO+DREADD mice only at 21 DPI. These results indicate that *RhoA;Pten* deletion in CSNs alone has limited promotion of forelimb motor recovery, however, the addition of excitatory DREADD significantly enhanced forelimb motor recovery after SCI.

**Figure 5.**
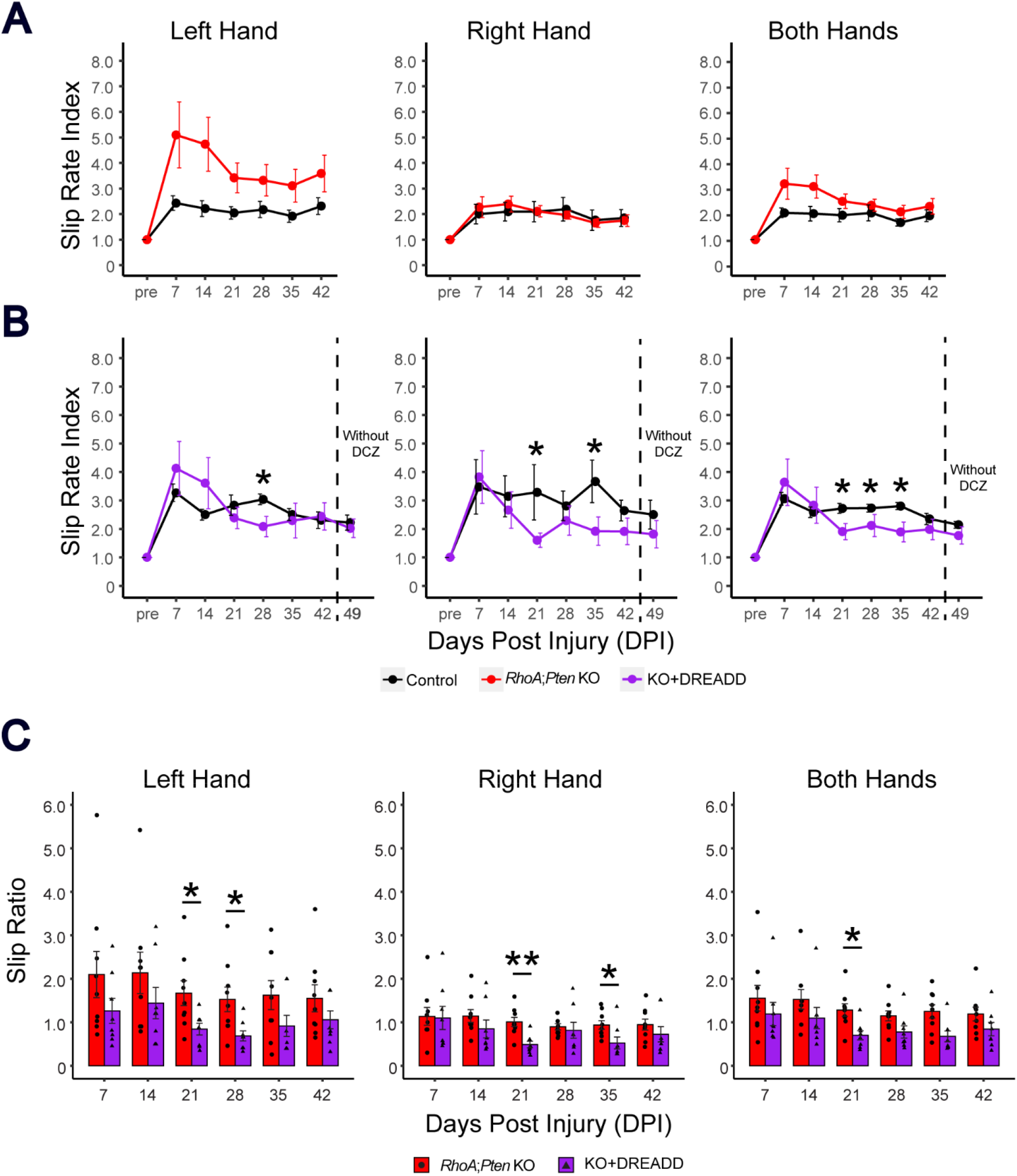
RhoA;Pten KO+DREADD enhances early forelimb motor recovery after SCI in the grid walking test. (A-B) Forelimb slip rate index of left, right and both hands in KO control mice (n = 7), RhoA;Pten KO mice (n = 9), KO+DREADD control mice (n = 7) and KO+DREADD mice (n = 8). Dotted lines indicate when DCZ administration was stopped. Wilcoxon rank sum exact test, *p<0.05. (C) Improvement rates of RhoA;Pten KO and KO+DREADD mice. Wilcoxon rank sum exact test, *p<0.05, **p<0.005.

### Combination of *RhoA;Pten* deletion and excitatory DREADD promotes axon sprouting after SCI

Activation of CSNs by excitatory DREADD at 21 DPI was evaluated by examining cFos expression in KO+DREADD mice and their controls (Fig. 7A). mCherry+ CSNs in KO+DREADD mice showed greater cFos expression than CSNs in controls, indicating the excitatory DREADD was induced successfully.

**Figure 6.**
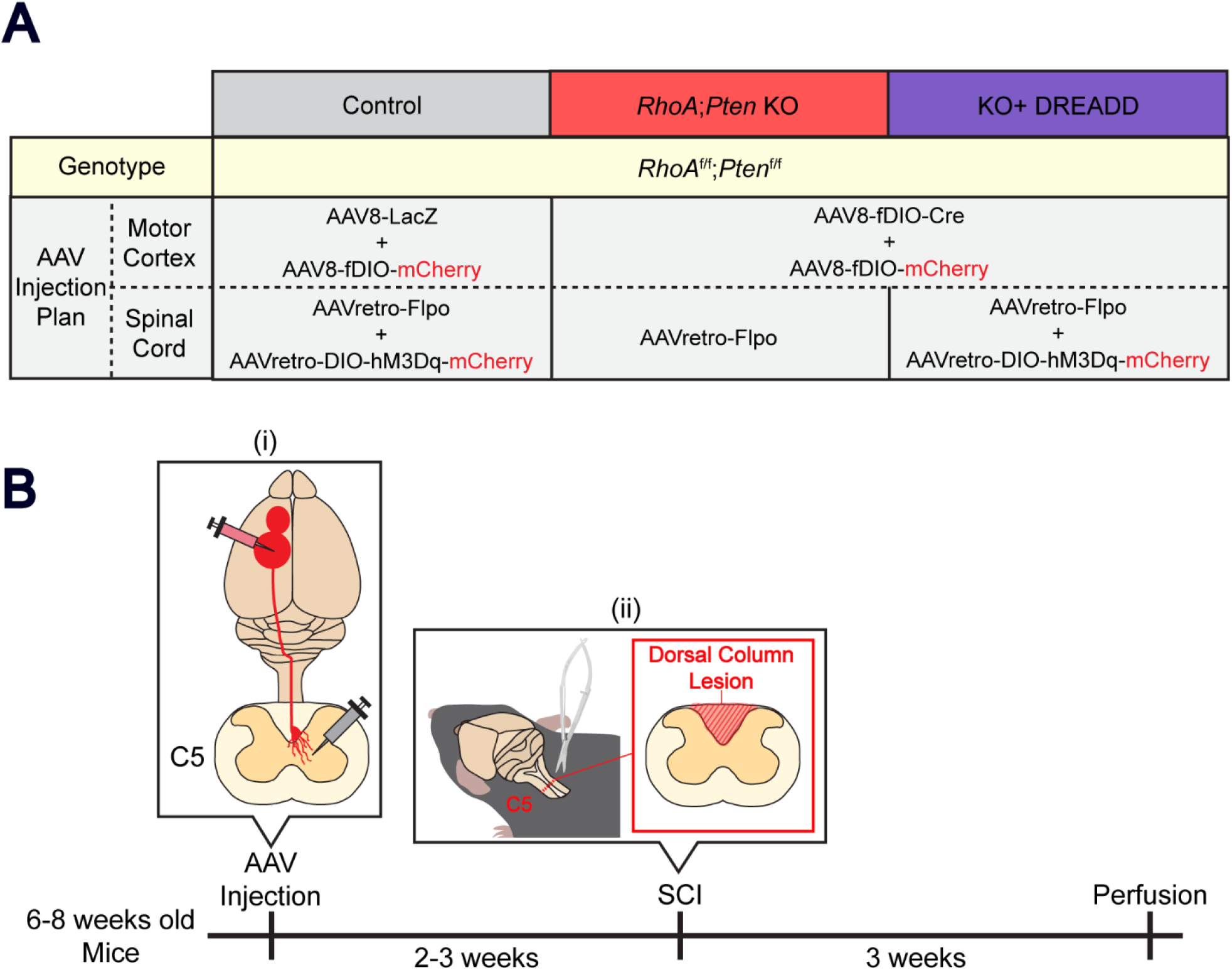
Experimental procedure of axon sprouting (A) AAV injection strategy. RhoA;Pten KO mice and KO+DREADD mice were generated by injecting AAVs to induce RhoA and Pten KO, hM3Dq, and mCherry expression in CST axons. Control mice were only injected with AAVs for mCherry expression. (B) Experimental schematic and timeline. (i) AAVs were injected unilaterally into the forelimb area of the sensorimotor cortex and the C5 level of the spinal cord in 6-8 week old RhoAf/f;Ptenf/f mice. (ii) The C5 dorsal column lesion was performed 2-3 weeks after AAV injections followed by perfusion at 3 weeks post-injury.

**Figure 7.**
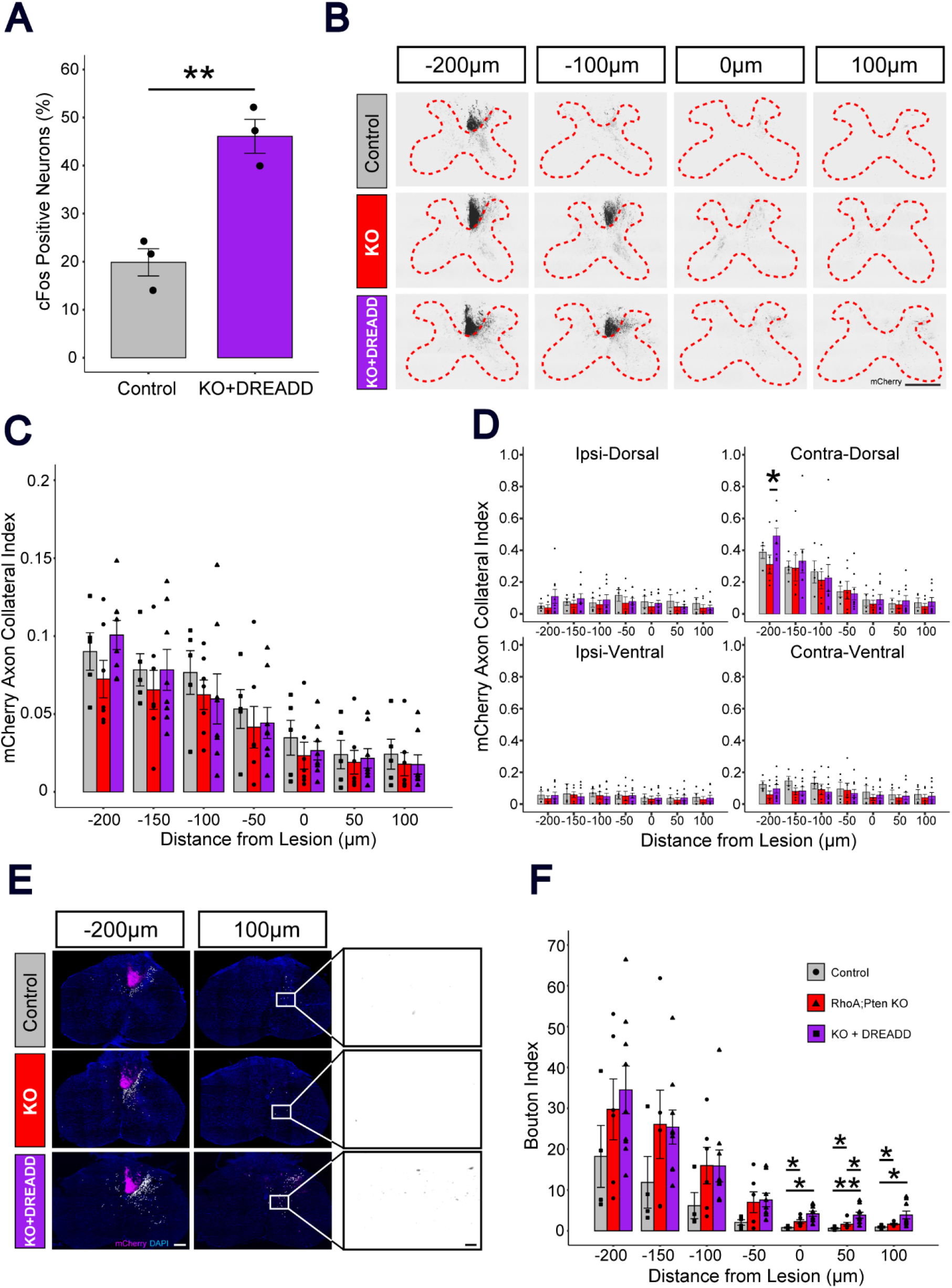
RhoA;Pten KO+DREADD promotes sprouting of CST fibers after SCI in the area rostral to the lesion. (A) Quantitative analysis of cFos+ mCherry neurons for control (n = 3) and KO+DREADD mice (n = 3). Unpaired t-test, **p<0.005. (B) Representative images of mCherry+ CST axons in the cervical spinal cord in control, RhoA;Pten KO and KO+DREADD mice at 21 DPI. Dotted red lines indicate the border between the white and grey matter. mCherry (black). Scale bar, 500 µm. (C) Quantification of CST axon collaterals in the grey matter in control (n = 4), RhoA;Pten KO (n = 4) and KO+DREADD mice (n = 7). Kruskal Wallis rank sum test followed by Wilcoxon rank sum exact test. (D) Axon collateral index in each section of the grey matter in control (n = 4), RhoA;Pten KO (n = 4) and KO+DREADD mice (n = 7). Contralateral-Dorsal (Top left), Ipsilateral-Dorsal (Top right), Contralateral-Ventral (Bottom left), Ipsilateral-Ventral (Bottom right). Kruskal Wallis rank sum test followed by Wilcoxon rank sum exact test. (E) Representative images of presynaptic boutons within the grey matter in control, RhoA;Pten KO and KO+DREADD mice at 21 DPI. White dots indicate the individual boutons in the grey matter. Left, mCherry (magenta) and DAPI (blue); right, mCherry (black). Scale bar, 200 µm (left) and 20 µm (right). Quantification of boutons in the grey matter in control (n = 4), RhoA;Pten KO (n = 4) and KO+DREADD mice (n = 7). Kruskal Wallis rank sum test followed by Wilcoxon rank sum exact test, *p<0.05.

We next examined axon collaterals in the spinal grey matter (Fig. 7B). In KO+DREADD mice, axon collaterals were more abundant at 200, 150, and 100 µm from the lesion site, but the differences compared to the KO mice and the control were not significant (Fig. 7C). In several cases, the axon collaterals in the spinal grey matter of KO+DREADD mice showed higher intensity at distances 200, 150, and 100 µm from the lesion site on the contralateral-dorsal side of the spinal grey matter, while other sections showed low intensities across all treatment groups (Fig. 7D). The difference at 200 µm was significant, indicating that *RhoA;Pten* KO combined with DREADD promotes axon sprouting after SCI.

Lastly, we evaluated the presynaptic connections in CSN axons by assessing the number of boutons in the spinal grey matter. The bouton indices, the amount of presynaptic connection by CSNs, in the KO and KO+DREADD mice were significantly higher than the control at the lesion and caudal of the lesion (Fig. 7E-F). Moreover, KO+DREADD mice showed significantly higher value than KO mice at 50 µm caudal to the lesion, indicating that addition of DREADD promotes the formation of presynaptic connections in the grey matter after SCI.

## DISCUSSION

A previous study showed that *RhoA*;*Pten* co-deletion prevents axon dieback after SCI but was insufficient to regain motor function^12^. Therefore, other strategies such as neuronal stimulation may need to be combined for motor recovery. Previous studies show that axon sprouting is promoted by neuronal stimulation, and the stimulation leads to recovery of motor function^13–16^.

Therefore, neuronal stimulation might enhance the axon growth/sprouting in *RhoA*;*Pten* co-deleted axons. In our present study, we combined genetic deletion of *RhoA* and *Pten* in CSNs with chemogenetic stimulation of CSNs and examined levels of axon regrowth and forelimb motor function recovery after cervical SCI. Our results in mice revealed that the additive effects of genetic deletion and CSN stimulation via excitatory DREADDs limited axon dieback and promoted structural connectivity of CSN axons at and below the site of SCI. Furthermore, this combinatorial treatment accelerated the recovery of skilled locomotor function after SCI.

An earlier study by our group showed that mice lacking *RhoA* and *Pten* exhibited less axon dieback than controls but did not promote axon regrowth through the lesion after SCI in the lumbar region. Hindlimb motor function was also not restored. Similarly, in our current study focused on cervical SCIs, we found that genetic deletion of *RhoA* and *Pten* as a singular strategy did not induce motor recovery in mice. However, when excitatory DREADDs that stimulate CSNs were combined with *RhoA; Pten*-deletion, axon dieback was reduced to a greater extent than mice that had only undergone *RhoA;Pten* deletion. Although the excitatory DREADDs and *RhoA;Pten* deletion promoted regrowth of injured CS axons, the axons did not extend across the lesion site and the motor function did not fully recover. Bridging the lesion and prolonging an immature state of neurons with neural stem cell (NSC) grafts ^26–28^, or neutralizing extracellular inhibitors at the lesion site ^27, 29, 30^, may promote greater CS regeneration in the spinal cord. These additional treatments may be required to achieve a greater degree of functional recovery.

Following dorsal hemisection injuries in which the CST is transected, mice lacking *RhoA* and *Pten* exhibit less axon dieback, but are still unable to recover hindlimb motor functions^12^. The effects on forelimb motor function were not reported. Given the critical role that the CST plays in voluntary skilled behaviors in the forelimb, we examined forelimb motor control in *RhoA;Pten* KO and *RhoA;Pten* KO+DREADD mice in a grid walking test^23, 31^. The KO+DREADD mice showed faster motor function recovery from 21 DPI compared to control. Although this combinatorial approach promoted early motor recovery after SCI, full motor function was never achieved.

Complete motor recovery may have been hindered for several reasons. First, the limited axon regrowth observed following SCI may be insufficient for regaining pre-injury levels of motor control. Other strategies (e.g., NSC grafts^26–28^, rehabilitation^32, 33^, chondroitinase treatment with peripheral nerve grafts^34, 35^) may need to be incorporated to further enhance motor recovery. A second possibility is that other circuits in addition to the CST, such as ascending sensory fibers, may also need restoration to regain full functionality. Finally, continuous administration of hM3Dq may result in its downregulation which would suppress the activation of CSNs^36^. Though complete motor recovery was not achieved, this is the first time even partial restoration of skilled movements has been demonstrated following SCI.

To determine how this partial motor recovery may occur, we examined bouton formation as an indicator of axon growth in *RhoA*;*Pten* KO and *RhoA*;*Pten* KO + DREADD mice. A prior study showed that *RhoA;Pten* deletion promotes axon sprouting after SCI at lumbar levels ^12^. In the present study, however, axon sprouting was absent in *RhoA;Pten* KO mice after cervical SCI, suggesting that circuit rewiring mechanisms in the cervical and lumbar spinal regions may differ.

The addition of excitatory DREADDs to stimulate CSNs in *RhoA;Pten* KO mice caused an increase in axonal boutons proximal and caudal to the cervical spinal cord lesion. Several studies have shown that excitatory DREADDs can promote presynaptic connections, while inhibitory DREADDs decrease the total number of boutons following SCI ^37, 38^. This suggests that CSN stimulation may augment the functional connectivity of therapeutic interventions that support axon outgrowth in individuals suffering from SCI.

Taken together, our results suggest that the effects of *RhoA*;*Pten* deletion combined with neuronal activation of CSNs is a promising therapeutic approach to promote axon regrowth and motor recovery after SCI. The recovery gains in motor function observed after SCI leave open the possibility that other multi-pronged approaches may yield even greater degrees of motor recovery. Future treatment paradigms may involve a mix-and-match model tailored to the type and location of spinal cord injury experienced by a person.

## ACKNOWLEDGMENTS

This work was supported by the Structural and Functional Imaging Core at Burke Neurological Institute, the New York State Spinal Cord Injury Research Board Grant C35599GG, and National Institutes of Health S10 Shared Instrumentation Grant OD028547-01. Y.Y. was supported by National Institute of Neurological Disorders and Stroke Grants NS100772, NS115963, NS119508, and NS093002. We thank I. Pavlova (Burke Neurological Institute) for technical advice of confocal images; Y. Zheng (Cincinnati Children’s Hospital Medical Center) for providing mice; and E. Hollis II and K. Friel (Burke Neurological Institute) for reading the manuscript.

## AUTHOR CONTRIBUTIONS

Conceptualization, H.T., I.F., and Y.Y.; methodology, H.T., I.F., and Y.Y.; data curation, H.T.; investigation, H.T.; formal analysis, H.T.; validation, H.T.; resources, H.T., I.F., and Y.Y.; writing – original draft, H.T.; review and editing, I.F. and Y.Y.; supervision, Y.Y.; project administration, Y.Y.; funding acquisition, Y.Y. All authors have read, edited, and approved the final manuscript.

## MATERIALS AND METHODS

### Animals

*RhoA^f/f^* ^17–19^ and *Pten^f/f^* (The Jackson laboratory) ^20^ mice were crossed to create *RhoA^f/f^;Pten^f/f^* mice maintained on C57BL/6 background. Mice were isolated in individual cages in a pathogen-free environment under a 12-hour light/dark cycle and fed commercial pellets and water ad libitum.

Procedures were performed in accordance with protocols approved by the Weill Cornell Medicine Institutional Animal Care and Use Committee.

### Adeno-Associated Viruses (AAVs)

The following AAVs were purchased from Addgene and used in experiments: AAV8-Ef1a-fDIO-Cre (AAV8-fDIO-Cre) [2.1 × 10^13^ GC/ml], AAV8-Ef1a-fDIO-mCherry (AAV8-fDIO-mCherry) [1.8 × 10^13^ GC/ml], AAV8-CMV-LacZ (AAV8-LacZ) [1.7 × 10^13^ GC/ml], AAV1-hsyn-Cre (AAV1-Cre) [1.9 × 10^13^ GC/ml], AAV1-hsyn-GFP (AAV1-GFP) [1.0 × 10^12^ GC/ml], AAVretro-Ef1a-Flpo (AAVretro-Flpo) [1.6 × 10^13^ GC/ml], AAVretro-hsyn-DIO-hM3Dq-mCherry (AAVretro-DIO-hM3Dq-mCherry) [1.9 × 10^1^^3^ GC/ml].

### Surgeries

AAV injections

Mice were anesthetized with isoflurane and placed in a stereotaxic frame (Stoelting, 51730D). For brain injections, the scalp was incised (incision size of 10 × 10 mm^2^), and holes were made at the corresponding sites of AAV injections of the skull using a round stainless-steel drill (11 0.5 mm). AAVs were injected into the RFA and CFA of the sensorimotor cortex (depth of 0.5 mm; coordinates, 1.8 mm anterior, 1.2 mm lateral to the bregma, 0.6 mm posterior, 1.8 mm lateral to the bregma ^21^, 0.2µl/virus/site) using a Nanoject III (Drummond scientific company, 3-000-207) tipped with a glass micropipette. For spinal cord AAV injections, skin and muscles on cervical region of the neck were incised, exposing the C4 vertebrae. AAVs were injected into the C5 region of the spinal cord (depth of 0.5 mm and 1.0 mm; coordinates, 0.5 mm anterior, 0.5 mm lateral to the center of C4 vertebrae, 0.5 mm posterior, 0.5 mm lateral to the center of C4 vertebrae, 0.15µl/virus/site) using the Nanoject III.

Spinal Cord Injury (SCI)

Dorsal column lesions were made using as previously described ^22^ . Briefly, after anesthetized with isoflurane, a laminectomy at the C4 vertebrae was performed to expose the spinal cord. A dorsal column lesion (depth of 1.0 mm) was made at the C5 vertebrae using Vannas spring scissors (2.5 mm cutting edge) to sever the dorsal CST.

### Analgesia

Prior to skin incision, a mixture of 2% lidocaine and 0.5% bupivacaine was administered subcutaneously with 30 G syringe to the incision site as a local analgesic. Buprenorphine (0.5 mg/kg) was administered subcutaneously with 30 G syringe as a postoperative analgesic immediately following surgery and twice per day thereafter for 3 days. Meloxicam (0.2 mg/kg) was additionally administered subcutaneously following AAV injections.

### Western blots

Dissected brains were sectioned using a mouse coronal brain slicer (Kent scientific, rbma-200c). Areas of the brain in which GFP fluorescence was detected were collected (Fig. 2A) and homogenized in lysis buffer (50 mM Tris-HCl, pH 8.0, containing 150 mM NaCl, 1% NP-40, 0.5% sodium deoxycholate, 0.1% SDS, and protease inhibitor cocktail (abcam, ab271306)). After centrifugation at 10,000 × *g* for 10 min at 4°C, the protein concentration was adjusted to 1 mg/ml, then proteins were separated by SDS-PAGE and transferred to a PVDF membrane (BIO-RAD, 162-0177). The membrane was blocked with 5% skim milk in PBS containing 0.05% Tween 20 and then incubated with rabbit anti-RhoA (1:1000, Cell Signaling Technology, 67B9), rabbit anti-Pten (1:1000, Cell Signaling Technology, 9188S), or mouse Tuj1 (1:1000, BioLegend, MMS- 435P) overnight at 4°C. After washing, the membrane was incubated with anti-rabbit IRDye 680RD (LI-COR, 925-68072) and anti-mouse IRDye 800 (LI-COR, 925-32213). An Odyssey Clx Imager (LI-COR) was used to detect and quantify antibody-bound proteins.

### Histology

Dissected brains and spinal cords were post-fixed in 4% PFA overnight. The tissues were then cryopreserved at 4 °C in 30% sucrose in PBS overnight and then embedded in Tissue-Tek optimal cutting temperature compound (Sakura Finetek). Embedded samples were sliced using a cryostat into 50 µm-thick sections and floated on PBS. The sections were then blocked with 1% bovine serum albumin in PBS for 1 hour before incubating with rat anti-GFAP (Thermo Fisher, 13-0300) or rabbit anti-cFos (Cell Signaling Technology, 2250S) antibodies overnight at 4°C. After washing with 0.1% Tween 20/PBS, the sections were incubated with Alexa Fluor 488 anti-rat (Invitrogen, A21208) or anti-rabbit IgG antibodies (Invitrogen, A21206). The sections were then washed with PBS and mounted onto glass slides (fisherbrand, 1255015). 200 µl of mounting media (VectorLabs, H10000-10) was applied directly onto the sections before covering with coverslips (VWR, 48393- 251). The slide-mounted sections were scanned using a confocal microscope (Nikon A1R HD25).

### Quantification of activated CSNs

To quantify cFos expression in CS neurons, coronal sections of the brain that exhibited mCherry^+^ CSNs were selected. Using ImageJ software, mCherry^+^ CSNs were first detected from confocal images to determine the region of interest (ROI) from the 16-bit red channel image. The intensity of cFos (fluorescently stained with Alexa Fluor 488) within the ROI was detected in the green channel image and the percentage of mCherry and cFos double-positive neurons was quantified.

### Quantification of CST dieback

CST dieback analysis was carried out following established quantification methods with some modifications ^12^. To quantify the dieback and regeneration of CST axons, the mCherry intensity within the CST fibers in the dorsal column were measured in sagittal section of the spinal cord by ImageJ software. The center of the lesion was set as 0 µm, and measurements were taken at 100 µm intervals along the rostral-caudal axis. To normalize the differences between individual animals, the mCherry intensity at each distance was divided by the intensity of mCherry in the rostral 1000 µm, and this value was defined as the axon index. To assess the suppressed dieback relative to control mice, the mCherry axon index of the *RhoA;Pten* KO mice and the KO+DREADD mice were divided by the mean index of the control mice, and this calculated value was defined as the axon dieback suppression ratio.

### Quantification of axon collateral projections

Axon collateral projection analysis was carried out following established quantification methods with some modifications ^12^. Transverse sections of the spinal cord at 0, 50, 100, 150, and 200 µm rostral, and 50 and 100 µm caudal to the lesion, were chosen to quantify axon collaterals. The mCherry intensity of CST fibers in the grey matter were measured in each section by ImageJ software. Since the C5-specific mCherry label does not stably show mCherry distal from the lesion and made it difficult to normalize individual differences in the control mice using mCherry intensity in the grey matter, the intensity in each section was divided by the intensity in the dorsal funiculus, 1000 µm rostral from the lesion. This value was defined as the mCherry axon collateral index.

### Quantification of presynaptic boutons

The total number of mCherry^+^ boutons was quantified by Imaris AI microscopy image analysis software. Boutons with diameters greater than 5 µm were selected and counted from the transverse sections of the spinal cord at 0, 50, 100, 150, and 200 µm rostral, and 50 and 100 µm caudal to the lesion site. The number of boutons was normalized by dividing by the mCherry intensity in the dorsal funiculus 1000 µm rostral to the lesion and the value was defined as the bouton index.

### Behavioral analyses

A grid-walking test using an elevated square wire grid (20 × 20 cm^2^, with each grid cell 1.2 × 1.2 cm^2^) was used to evaluate the effects of SCI on the skilled behaviors of treated and control mice. A custom-made acrylic transparent tube (1125 cm diameter) was placed on the grid to prevent mice from walking on the edge of the grid. Limb slips were detected during playbacks of video recordings (30 fps) of traversal attempts. A mirror was placed under the grid at an angle of 45° and the reflection was recorded with another video camera to identify which limb slipped off the grid.

Mice were subjected to the grid-walking test prior to SCI (pre-injury), then 7, 14, 21, 28, 35, 42, and 49 days post-injury (DPI). Mice were allowed to walk on the grid for 3 min and foot-slips of the left and right forelimbs were counted during the first 50 forelimb steps. A slip was scored when the paw completely missed the grid and the limb fell between the wires, or when the paw was correctly placed on the grid but slipped off during the weight-bearing phase ^23^. The percentage of slips was calculated as: the number of slips divided by the first 50 steps per trial × 100. To normalize differences between animals, the slip rate at each DPI was divided by the pre-injury rate, and this value was defined as the slip rate index. To show deterioration from controls, the slip rate index of *RhoA;Pten* KO mice and KO+DREADD mice were divided by the mean index of the control mice, and this value was defined as the slip ratio.

### Experimental Design

For *RhoA;Pten* KO confirmation, 6 weeks old *RhoA^f/f^;Pten^f/f^* mice were injected with AAV1-GFP or AAV1-Cre in the rostral and caudal forelimb areas (RFA and CFA) of the sensorimotor cortex (Fig. 2A). Two weeks later, animals were euthanized by CO_2_ exposure, brains were dissected and processed for western blotting.

To assess the level of axon regrowth and motor behavior in the forelimb following *RhoA;Pten* KO and excitatory DREADD, 6-8 week-old *RhoA^f/f^;Pten^f/f^*mice were first injected with AAVretro-Flpo to induce mCherry or Cre expression in the CST (Fig. 1C). RhoA;Pten KO or hM3Dq were induced by Cre in two mouse groups (the RhoA;Pten KO mice and the KO+DREADD mice), but KO and hM3Dq were absent in control mice due to the lack of Cre (Fig. 1A-B). After 2-3 weeks, SCI was made in C5 spinal cord and weekly grid walk test starting from one week after SCI was performed. Deschloroclozapine (DCZ, MCE, HY-42110) was administered intraperitoneally with 30G syringe to AAVretro-DIO-hM3Dq-mCherryinjected mice twice a day at a concentration of 100 µg/kg^24, 25^ from the day after injury until 42 days post-injury (DPI). For the days when mice were subjected to the weekly grid walk test, DCZ was administered 1 hour before and 1 hour after the test. To confirm whether hM3Dq is affecting behavior in the KO+DREADD and control mice, an additional week of grid-walking tests were performed without DCZ. After the final grid-walking test, the *RhoA;Pten* KO mice and their controls were transcardially perfused with 4% PFA in phosphate buffer, while the KO+DREADD mice and their controls were administered with DCZ and perfused 2-3 hours later. The brain and spinal cord were dissected and processed for immunohistochemistry.

To quantify axon collateral projections, 6-8 week-old *RhoA^f/f^;Pten^f/f^* mice (comprised of controls, *RhoA;Pten* KO and KO+DREADD mice) were injected with AAVs unilaterally according to the injection plans (Fig. 6A-B). SCI was made in the spinal cord at cervical level 5 (C5) 2 weeks after AAV injections, mice were perfused at 21 DPI, and brains and spinal cords were dissected and processed for immunohistochemical analyses (Fig. 6C).

### Statistical analyses

The numbers of animals used in each experiment are reported in the Results section and the figure legends. Quantitative data are represented as the mean ± SEM. Statistical analyses were performed using R software. Differences among groups were statistically analyzed using a two-tailed unpaired t-test or Wilcoxon rank sum exact test. A *p*-value of <0.05 was considered statistically significant.

